# Functional connectivity, tissue microstructure and T2 at 11.1 Tesla distinguishes neuroadaptive differences in two traumatic brain injury models in rats: A Translational Outcomes Project in NeuroTrauma (TOP-NT) UG3 phase study

**DOI:** 10.1101/2023.12.10.570975

**Authors:** Rohan S. Kommireddy, Shray Mehra, Marjory Pompilus, Rawad Daniel Arja, Tian Zhu, Zhihui Yang, Yueqiang Fu, Jiepei Zhu, Firas Kobeissy, Kevin K.W. Wang, Marcelo Febo

## Abstract

Traumatic brain injuries (TBIs), particularly contusive types, are associated with disruptions in neuronal communication due to focal and diffuse axonal injury, as well as alterations in the neuronal chemical environment. These changes can negatively impact neuronal networks beyond the primary injury site. In this Translational Outcomes Project in NeuroTrauma (TOP-NT) UG3 phase study, we sought to use multimodal neuroimaging biomarker approach to assess functional connectivity and brain tissue microstructure, along with T2 relaxometry, in two experimental rat models of TBI: controlled cortical impact (CCI) and lateral fluid percussive injury (LFPI). Rats underwent imaging using an 11.1 Tesla scanner at 2 and 30 days post-injury. Naïve controls were scanned once to establish baseline comparisons for both TBI groups. Imaging modalities included functional magnetic resonance imaging (fMRI), diffusion-weighted imaging (DWI), and multi-echo T2 imaging. fMRI data were analyzed to evaluate functional connectivity across lateral and medial regions of interest (ROIs) in the cortical mantle, hippocampus, and dorsal striatum. DWI scans were used to generate maps of fractional anisotropy (FA) and mean, axial, and radial diffusivities (MD, AD, RD), focusing on cortical and white matter (WM) regions near the injury epicenter. Our findings revealed significantly increased contralateral intra-cortical connectivity at 2 days post-injury in both CCI and LFPI models, localized to similar cortical areas. This increased connectivity persisted at day 30 in the CCI model but not in LFPI. Changes in WM and cortical FA and diffusivities were observed in both models, with WM alterations predominating in CCI and cortical changes being more pronounced in LFPI. These results highlight the utility of multimodal MR imaging for characterizing distinct injury mechanisms in contusive and skull-penetrating TBI models.

## Introduction

Traumatic brain injury (TBI) affects millions worldwide and is associated with mortality rates exceeding 30%^1^. While mild-to-moderate repetitive closed head injuries are the most common types of TBI, focal contusions and skull-penetrating injuries caused by forceful blows, jolts, or impacts to the head, neck, or spinal cord often result in severe outcomes. These injuries can lead to lifelong sensorimotor, emotional, and cognitive impairments, along with permanent disruptions in brain function and structure. The profound emotional and financial burdens on patients and caregivers underscore the need for improved therapeutic strategies. The heterogeneity of brain damage in severe TBI complicates clinical management, as its pathophysiology involves diverse and complex cellular and molecular mechanisms extending beyond the primary injury site.

Secondary injuries, which develop after the initial trauma, can include aberrant neuroimmune responses, diffuse axonal injury (DAI), white matter (WM) damage, blood-brain barrier (BBB) disruption, microvascular bleeding, intracranial hematomas, cerebral and subdural hemorrhages, edema, epileptiform activity, and other maladaptive central nervous system (CNS) pathologies^2^. Effective clinical management of TBI hinges on early intervention to mitigate the progression of these secondary injuries^3^. Despite promising neuroprotective effects observed in experimental drugs using preclinical TBI models, none have shown efficacy in clinical trials^4^. This gap in translational success is partly due to a lack of minimally invasive, in vivo CNS biomarkers capable of bridging preclinical findings with clinical applications^4^.

Functional neuroimaging has the potential to provide translational data that link mechanisms of disease progression to behavioral recovery in TBI, especially when combined with other quantitative magnetic resonance imaging (MRI) modalities. For instance, studies of mild-to-severe TBI have shown stronger deactivation of default mode network (DMN) regions during cognitive tasks and a negative correlation between posterior cingulate connectivity and mean diffusivity (MD) in the splenium^5^. Similarly, mild TBI has been associated with reduced connectivity in these regions^6^, although hyperconnectivity within the DMN has also been reported^7^ and linked to metacognitive recovery in moderate-to-severe TBI^8^. Temporal shifts in functional connectivity, characterized by both increases and decreases, have been observed in limbic, sensorimotor, and associative cortical areas following moderate-to-severe^9^ and mild^10^ TBI. For example, sports-related concussions have been associated with decreased long-distance and increased local functional connectivity^11^.

Despite the variability in primary and secondary damage caused by TBI, common patterns emerge, such as temporally evolving reductions and increases in functional connectivity across cortical subregions. However, it remains unclear whether these changes are restricted to specific cortical domains, such as cognitive, motor, sensory, or association cortices, over time. Animal models replicating mild concussions from sports injuries or automobile accidents are essential for understanding prevalent forms of TBI in humans^12,13^. While these models provide valuable insights into the diffuse and heterogeneous pathology seen in clinical TBI, they have shown limited ability to elucidate the fundamental mechanisms driving cortical reorganization observed with contusions and penetrating injuries.

Preclinical studies using controlled cortical impact (CCI) and lateral fluid percussive injury (LFPI) models consistently demonstrate functional reorganization of the cortex both near and distant from the injury epicenter^14^. This large-scale network reorganization persists for 30 to 60 days post-injury^15,16^ and is thought to involve excitatory-inhibitory rebalancing^17^. Cortical changes are likely accompanied by alterations in WM microstructure^14^ and chemical composition detectable through tissue T2 relaxometry^18^.

The objective of the present study was to assess functional connectivity, brain tissue microstructure, and T2 relaxometry in CCI and LFPI models to identify key similarities and differences between these two models of contusive and penetrating TBI.

## Materials and Methods

### Subjects

Sprague-Dawley rats (220-300g; N=92, including 34 female rats) were obtained from Charles River Laboratories (Raleigh, NC, USA.). Rats were housed in sex-matched pairs in a temperature and humidity-controlled vivarium with a 12 h light cycle (lights on at 0700 h) and food and water provided ad libitum. Rats were randomly assigned to one of three experimental conditions: controls (n=29, including 8 female rats), CCI (n=31, including 13 female rats), and LFPI (n=13, including 5 female rats). CCI and LFPI groups were scanned on day 2 post-injury and a subset of these were reimaged at thirty days post-TBI. Day 30 CCI groups included. The latter subset included 11 CCI (8 females) and 8 male LFPI rats. The number of rats that received diffusion weighted imaging (DWI) and T2 relaxometry were lower than for fMRI due to the removal of scans. The number of rats receiving DWI and T2 scans are provided in corresponding figure legends. All procedures received prior approval from the Institutional Animal Care and Use Committee of the University of Florida and followed all applicable NIH guidelines.

### Controlled Cortical Impact (CCI)

The CCI procedure was carried out using a Leica Impact One stereotactic impactor (Leica Biosystems Inc), as part of the translational outcomes project in neurotrauma (TOP-NT) UG3 phase study. Anesthetic levels were induced with 4% isoflurane gas mixed with 100% oxygen and maintained under 2% isoflurane for the rest of the procedure. The body temperature was regulated to 37°C using a thermal pad while rats were prepared for surgery on a stereotaxic frame. A parasagittal craniectomy (center at anteroposterior [AP], -4.0 mm; L, +2.8 mm from lambda) 5 mm in diameter was performed to expose the brain and allow impactor tip access to the cortical surface (right side). The impactor had a 4-mm flat-face tip. CCI at a depth of 1.5 mm at 4lJm/s and a dwell time of 240ms was carried out. All injuries occurred in the right hemisphere. The surgical area was sutured, and recovery was monitored by tail pinch and righting reflexes. Control rats were naïve and received no sham surgical procedures.

### Lateral Fluid Percussion Injury (LFPI)

To model diffuse and focal open-head TBI and as part of the translational outcomes project in neurotrauma (TOP-NT), we used the rat LFPI model. Surgical preparation and maintenance were similar to that described for CCI, but used a pendulum-driven lateral impact via a fluid filled chamber delivering pressure pulse to a tube overlying an exposed area of the right cortex at the same coordinates used with the CCI procedure^19^. LFPI is a well-characterized model of TBI that captures clinically relevant deficits and pathologies^20–22^. LFPI also applies a localized, mechanical insult through a parasagittal craniectomy (right side). LFPI is administered via a pressurized fluid pulse (2.0 ±0.2atm). This injury produces moderated diffuse and focal injury on the right sensorimotor cortex and expansive and progressing white matter injury^23^ and neurobehavioral deficits^21,24^.

### Magnetic Resonance Imaging (MRI)

Images were collected on an 11.1 Tesla MRI scanner (Magnex Scientific Ltd., Oxford, UK) with a Resonance Research Inc. gradient set (RRI BFG-240/120-S6, maximum gradient strength of 1000 mT/m at 325 Amps and a 200 µs risetime; RRI, Billerica, MA) and controlled by a Bruker Paravision 6.01 console (Bruker BioSpin, Billerica, MA). A custom-made 2.5 cm x 3.5 cm quadrature radiofrequency (RF) surface transmit/receive coil tuned to 470.7MHz (^1^H resonance) was used for B1 excitation and signal detection (RF engineering lab, Advanced Magnetic Resonance Imaging and Spectroscopy Facility, Gainesville, FL). Rats were scanned under a continuous flow of 1.5 % isoflurane (delivered at 0.1L/min mixed with medical-grade air containing 70% nitrogen and 30% oxygen). Respiratory rates were monitored continuously, and body temperature was maintained at 36-37°C using a warm water recirculation system (SA Instruments, Inc., New York).

A high-resolution T2 weighted anatomical scan was acquired with a functional magnetic resonance imaging (fMRI) scan, a diffusion weighted imaging (DWI) scan, and a multi-echo time image series for T2 mapping. The T2 weighted Turbo Rapid Acquisition with Refocused Echoes (TurboRARE) scans were acquired with the following parameters: effective echo time (TE) = 37 ms, repetition time (TR) = 5 seconds, RARE factor = 16, number of averages = 14, field of view (FOV) of 24 mm x 18 mm and 0.9 mm thick slice, and a data matrix of 256 x 192 and 25 interleaved ascending coronal (axial) slices covering the entire brain from the rostral-most extent of the anterior frontal cortical surface, caudally towards the upper brainstem and cerebellum. Functional images were collected using a single shot spin echo planar imaging (EPI) sequence with the following parameters: TE = 15 ms, TR = 2 seconds, 300 repetitions, FOV = 24 x 18 mm and 0.9 mm thick slice, and a data matrix of 64 x 48 with 25 interleaved ascending coronal slices in the same position and orientation as the corresponding anatomical. Ten dummy EPI scans were run prior to acquiring data under steady state conditions. Respiratory rates, isoflurane concentration, body temperature, lighting, and room conditions were kept constant across subjects. Diffusion images used a 4-shot spin echo EPI readout with TE=18ms, TR=4 seconds, 4 averages, FOV = 24 x 18 mm and 0.9 mm slices and a data matrix of 128 x 96 x 25, 30 directions with a b value = 900 s^2^/mm and 4 b = 0 scan, pulse duration = 3 ms, and gradient separation = 8 ms. Multi-slice multi-echo scans were collected with 10 TE’s from 6-60 ms in 6 ms steps per image volume, TR=2 seconds, 2 averages, FOV = 24 x 18 mm with 0.9 mm slices and a data matrix of 128 x 128 x 25.

### Functional MRI (fMRI) Processing

Image processing used Analysis of Functional NeuroImages (AFNI)^25^, FMRIB software library (FSL) version 6.0.5^26^, and Advanced Normalization Tools (ANTs)^27^ and manual mask drawing and editing functions in ITKSNAP^28^. Masks outlining the brains of rats on fMRI, DWI, and T2 anatomical scans were generated using *3dAutomask* in AFNI. These were used for brain extraction in various steps in the image processing workflows. The processing of fMRI scans included the following steps: (1) timeseries spike identification and removal using *3dDespike* in AFNI, (2) motion correction using *3dvolreg* in AFNI, (3) removal of slow signal drifts (<0.009Hz) using *3dTproject* in AFNI, (4) identification and suppression of nuisance signals using *fsl_regfilt* in FSL^29^ and (5) low-pass filtering (>0.12Hz) and spatial smoothing (0.6mm FWHM).

Preprocessed functional images were next co-registered to create a multisubject functional MRI template, using previously described methods^29^. The linear registration step used the following parameters: gradient step size=0.1, shrink factors= 4 x 2 x 1, smoothing factors= 2 x 1 x 0 voxels, matrix iterations=30 x 20 x 4, iteration limit = 3, a cross-correlation similarity metric, and rigid body followed by affine registration. The nonlinear registration step used similar parameters but with an iteration limit = 4 and a Greedy Symmetric Normalization (SyN) transformation model. Individual co-registration matrices were used to transform each subject’s fMRI scan to the functional template space. Whole brain statistical analyses were carried out in template space at the fMRI scan resolution. The multisubject template and statistical maps were then registered to an segmented atlas of the rat brain for 3D connectome visualizations^13^.

Group probabilistic independent components analysis (ICA) was used to evaluate functional connectivity network differences between control and TBI groups. Functional MRI voxel time series were thresholded, variance normalized, pre-whitened, and projected to a 20- and a 60-dimensional principal component subspace in separate analyses. The decomposed signals were optimized according to their non-Gaussian spatial source distributions using a fixed-point iterative approach ^30^. The final component maps were divided by the standard deviation of the residual noise and thresholded by fitting a mixture model to the histogram of intensity values ^31^. We used a 0.6 mixture model threshold for the 20 component ICA and 0.9 for the 60 component ICA, both with full temporal concatenation. The 20-dimensional ICA approach enabled detection of previously established default mode like somatomotor, anterior cingulate, and striatal networks in isoflurane sedated rats ^32^ (**Supplemental Fig. 1**). A dual regression approach was used to project spatial components back to each subjects native space and produce subject-specific timeseries for each spatial component ^33^. The subject-specific 20 component maps were statistically analyzed for *within-network* differences with 5000 permutation tests under a general linear model framework using FSL *randomise* ^34^. A statistical design matrix was created in FSL Glm for a one-way ANOVA with overall F-tests and specific post hoc contrasts between TBI groups and controls. Threshold free cluster enhancement (TFCE) and family wise error rate (FWER) correction were applied to all statistical p-value maps. Time series from the dual regression step were imported to MATLAB to construct network matrices. Pearson cross correlations were carried out and the resulting r coefficients were Fisher z-transformed prior to statistical comparisons. Matrices were analyzed using brain connectivity toolbox, as previously described ^29^. We focused our analysis on node-based network measures for graph densities of 10,15 and 20% and included assessments of network strength, assortativity, transitivity, path lengths and efficiency for weighted undirected 60^2^ matrices with Fisher’s z converted Pearson correlation coefficients. Formal descriptions of these graph theory measures, as applied to functional neuroimaging data in human subjects, is provided by Rubinov and Sporns^35^, and for rodents in our previous publications^15,29,36^. Fisher z statistics were rescaled from 0 to 1 prior to calculations of transitivity, a whole brain network variant of the clustering coefficient. We analyzed ROI node strengths at 15% graph density threshold. This threshold is based on prior functional MRI studies in rats imaged at 11.1 Tesla^37^ and is within the wide range of graph density thresholds described for various types of brain histological based connectome datasets for several species^38–42^. Whole brain statistical analysis was conducted with FSL permutation based *randomise* tool for ICA networks.

We used an approach recently published in Ontiveros-Ángel *et al*^43^ to determine structural differences between groups. Anatomical T2 scans were co-registered to create an anatomical template and assess structural differences by analyzing log normalized Jacobian nonlinear warping matrices. This provided additional quantitative assessments of brain-wide structural changes with CCI and LFPI at 2- and 30-days post-injury. Images were subject to the same pipeline described above for fMRI scans. Diffeomorphic matrices resulting from nonlinear warps in ANTs were log normalized using *CreateJacobianDeterminantImage* script in ANTs and analyzed using the *randomise* tool in FSL, with the same ANOVA model used in the analysis of ICA datasets.

### Diffusion Weighted Image (DWI) Processing

ITKSNAP was used to segment contralateral and ipsilateral cortical and white matter ROIs. Diffusion MRI scans were processed using tools available on FSL ^44^. DWI scans were first screened for volumes with low signal-to-noise or artefacts. Eddy correction was used for adjusting slight movements during image acquisition and gradient files rotated according to the motion correction vectors. After eddy correction, tensor element reconstruction and estimates of eigenvectors and eigenvalues were performed using weighted least squares regression on *dtifit* in FSL. This last step generated independent images of FA, mean, axial and radial diffusivities (MD, AD, and RD, respectively).

### T2 Mapping

T2 parametric maps were created using the MRI processor plugin for ImageJ^45,46^. The simplex minimization method was used for non-linear fitting of multi-TR images, using the equation S_TE_ = S_0_ * e^−TE/T2^. ITKSNAP was used to segment contralateral and ipsilateral cortical and white matter ROIs, similar to DWI scans. From T2 maps, the mean T2 is calculated per ROI and exported for statistical analysis.

### Statistical Analysis

Statistical analyses and data plotting were carried out using GraphPad Prism 9. Data were analyzed using repeated measures analysis of variance (ANOVA: group x density or ROI, significant p < 0 .05) with Tukey’s post hoc multiple comparison test.

## Results

### CCI produces persistent functional changes in contralateral brain networks with spill-over connectivity changes in bilateral regions

From the fMRI data, we identified the 20 ICA functional networks, which included previously reported retrosplenial, somatosensory, parietal, motor, striatal/insular networks (**Supplemental Fig. 1**). CCI increased functional connectivity within a contralateral parietal/somatosensory cortical network on day 2 post-injury (**Fig. 1**) (F_4,87_=4.3, p = 0.003; Tukey’s p = 0.003). The increased connectivity with the left (contralateral) parietal/somatosensory barrel field (S1BF) ICA component on day 2 was densely distributed with spill over across bilateral (contra and ipsilateral) cortical and subcortical areas, including visual, motor and striatal regions. This CCI-induced increase in contralateral S1BF network connectivity persisted through day 30 post-injury (F_4,87_=4.6, p = 0.002; Tukey’s p = 0.003). As observed on day 2, the brain regions showing increased connectivity with contralateral S1BF included both contra and ipsilateral regions (**Fig. 1B**). In addition, CCI increased functional connectivity with a midline hypothalamic network on day 30 (F_4,87_=7.0, p < 0.0001; Tukey’s p = 0.0003). The patterns of connectivity with this network were less distributed and included ipsilateral striatal, contralateral insular, contralateral ventral hippocampal and ipsilateral somatosensory cortex near the CCI epicenter.

**Figure 1.**
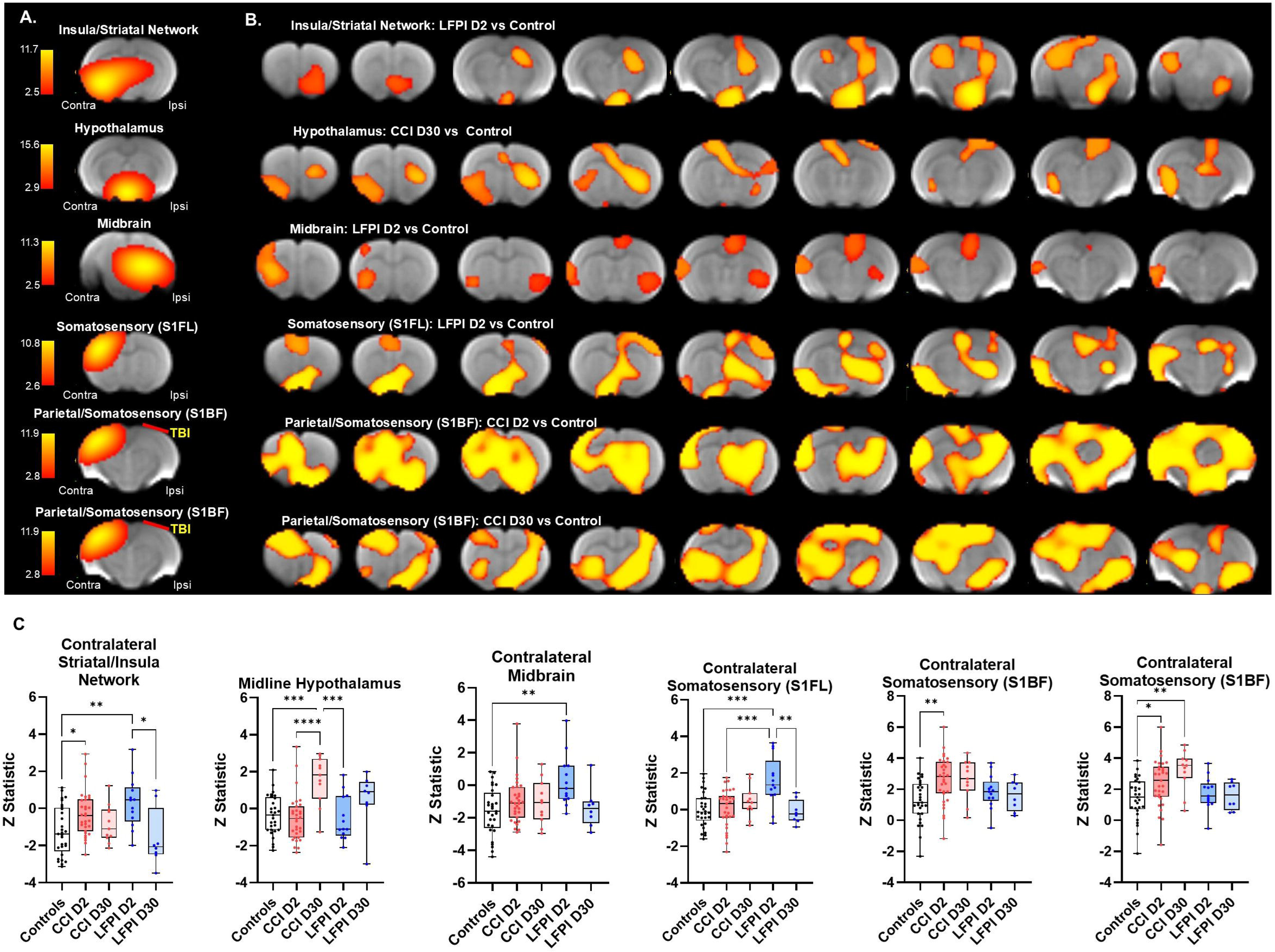
Independent components analysis (ICA) reveals cortical and subcortical networks distinctly affected by controlled cortical impact (CCI) and lateral fluid percussion injury (LFPI). A) ICA networks with significant differences between controls and TBI groups (scale bar color indicates a range of significant t-statistical values). B) Specific group comparisons within each network reveals brain areas functionally affected by CCI and LFPI on days 2 and 30 post-injury (p < 0.05, corrected). C) Functional connectivity differences in the same brain areas identified in B. One way analysis of variance (ANOVA) with Tukey’s multiple comparisons test (*p<0.05, **p<0.01, ***p0.001, ****p<0.0001). Data shown as median and range with overlaid individual data points.

In contrast, functional differences in response to LFPI were limited to day 2 post-injury. No effects of LFPI vs control were observed on day 30. FPI increased connectivity in a contralateral insular cortex/striatal network (F_4,87_=5.1, p = 0.001; Tukey’s p = 0.004), an ipsilateral midbrain network (F_4,87_=4.0, p = 0.005; Tukey’s p = 0.002) and a contralateral somatosensory network (F_4,87_=6.3, p = 0.0002; Tukey’s p = 0.0002) identified in the contralateral forelimb region (S1FL) (**Fig. 1**). Increased functional connectivity with the contralateral insula/striatum involved predominantly ipsilateral dorsal hippocampal and midline hypothalamic areas in LFPI rats compared to controls. Increased connectivity with the midbrain network involved contralateral anterior insular cortex and bilateral posterior insular cortex spill over and ipsilateral cingulate cortex. Differences in functional connectivity with the S1FL involved bilateral anterior cingulate, midline diagonal band region of the forebrain with spill over to ispilateral bed nucleus of stria terminalis (BNST) and ipsilateral dorsal striatum, contralateral olfactory cortex, ipsilateral dorsal hippocampus, and ipsilateral S1FL.

### CCI produces both transient and persistent changes in functional network topology that encompasses both ipsilateral and contralateral nodes

We used a set of ICA generated nodes to evaluate whole brain network and local node connectome measures on day 2 and 30 post-injury, in both CCI and LFPI rats. Connectome measures were assessed for graph densities of 10, 15, and 20%. The functional connectome measures reveal topological features of intrinsic fMRI signal covariations instead of fMRI signals preselected by a parcellation^47^. Consistent with our previous analysis^15^, connectome maps highlight greater node strength values on the contralateral compared to ipsilateral hemispheres in both CCI and LFPI rats on days 2 and 30 (**Fig. 2A**). Thus, despite ICA nodes being bilaterally distributed, their individual strength values emphasized a greater strength on the contralateral hemisphere relative to the ipsilateral site. Maps in **Fig. 2B** are for modularity, with node sizes indicating the module affiliation codes. The maps highlight a relatively stable and consistent rostral to caudal organization of modules across all groups. However, CCI day 2 maps show altered modular organization for nodes at the TBI epicenter (ipsilateral cortex) and spillover of modular nodes to the contralateral region.

**Figure 2.**
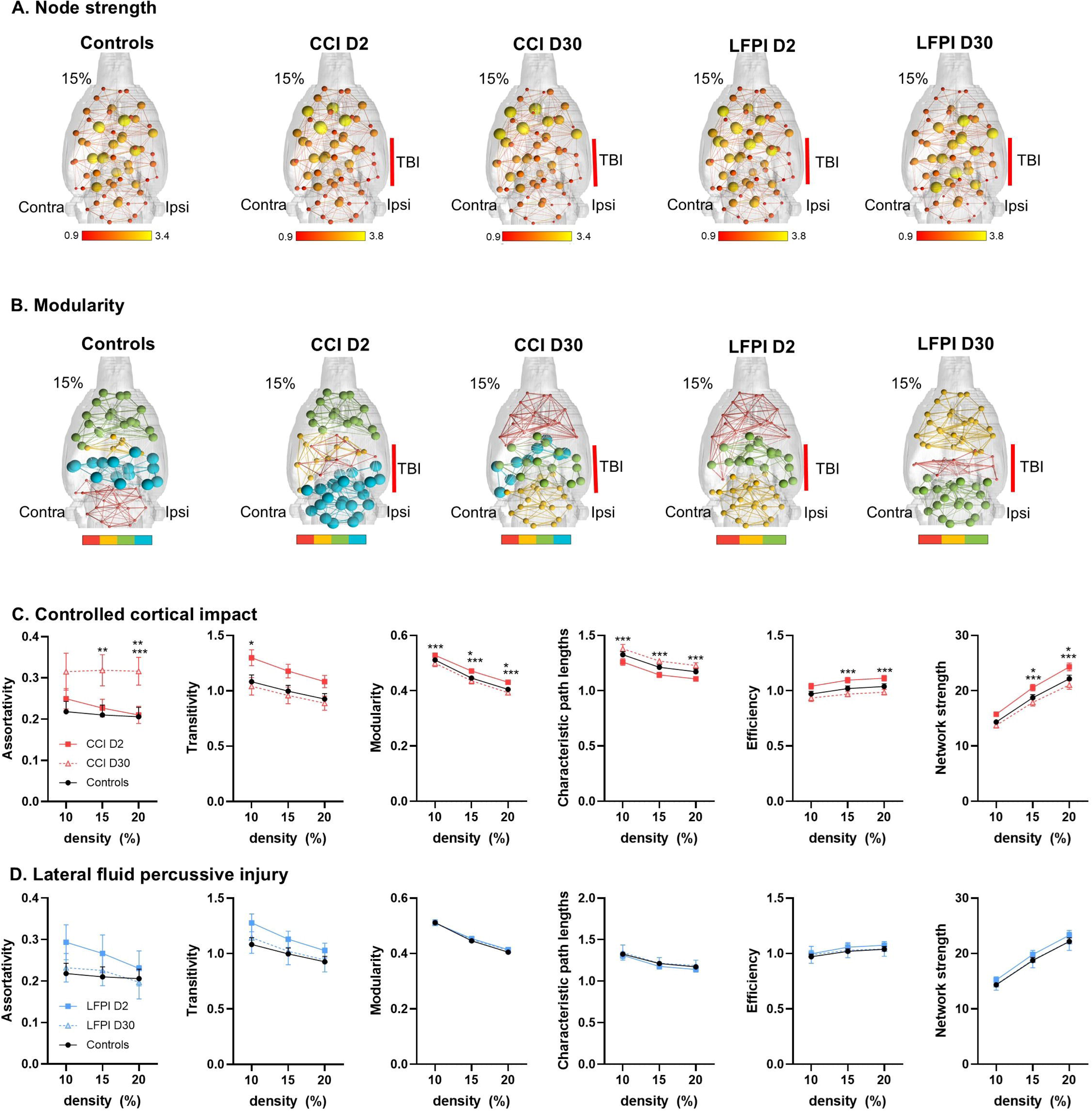
Functional network strength and modularity significantly differs with CCI but not LFPI. A) Node strength connectome maps for functional connectivity matrices averages within each group. Node sizes are node strength calculated for matrices at 15% graph density and lines connecting nodes are pairwise Pearson correlations. The scalebar color is for Fisher’s z transformed Pearson correlations. B) Modularity connectome maps for the same matrices in A. Node sizes and color are based on estimated module affiliations of each node. Lines are as in A but color-matched to each module. The scalebar below the maps indicate module affiliation. C) Network measures for CCI days 2, 30 and controls calculated at graph density thresholds of 10, 15, and 20%. D) Network measures for LFPI days 2, 30 and controls. Two-way ANOVA comparisons followed by Tukey’s multiple comparisons test (*day 2 vs control, ** day 30 vs control, *** day 30 vs day 2). Data shown as mean ± standard error. The approximate craniotomy/impact site for CCI and LFPI are shown on 3D connectome maps.

CCI altered whole brain network transitivity, modularity, path lengths, efficiency and network strength on day 2 that returned to control levels by day 30. This indicates that CCI results in a greater integration of node activity while increasing segregation or functional specialization across groups of nodes involving a heightened network efficiency. Conversely, assortativity differences, that were not present on day 2 post-injury, emerged on day 30 (**Fig. 2C**). This indicates that on day 30 the network topology of CCI rats involves an increased probability of strongly connected nodes with other strongly connected nodes and weaker nodes interacting with weak nodes. No differences in network measures were observed on day 2 and 30 following LFPI (**Fig. 2D**). It is important to point out that these network metrics are whole brain and do not specifically report on contralateral and ipsilateral changes.

We further analyzed node strength values at a graph density of 15% in CCI and LFPI relative to controls (**Fig. 3**). CCI produced effects on node strength with increases in several nodes on days 2 and 30 and mostly decreases in node strength in a subset of nodes on day 30 (group x ROI interaction: F_118,4012_ = 0.0007; group effect: F_2,68_ = 4.5, p = 0.01). CCI increased node strength in the midline anterior cingulate cortex (p = 0.002, CCI day 2 vs control), contralateral visual cortex (p < 0.0001, CCI day 2 vs control; p = 0.009, CCI day 30 vs control), contralateral S1BF (p = 0.01, CCI day 2 vs control), contralateral anterior insular cortex (p = 0.01, CCI day 2 vs control), contralateral motor cortex (p = 0.009, CCI day 2 vs control), contralateral hindlimb cortex (p = 0.005, CCI day 2 vs control), and contralateral auditory/temporal cortex (p= 0.001, CCI day 2 vs control; p = 0.007, CCI day 30 vs control). CCI reduced node strength in contralateral amygdala (p=0.02, CCI day 30 vs 2), ipsilateral visual cortex (p=0.03, CCI day 30 vs control), contralateral diagonal band (p=0.03, CCI day 30 vs day 2), ipsilateral laterodorsal thalamus (p = 0.03, CCI day 30 vs control), and ipsilateral BNST (p=0.008, CCI day 30 vs day 2).

**Figure 3.**
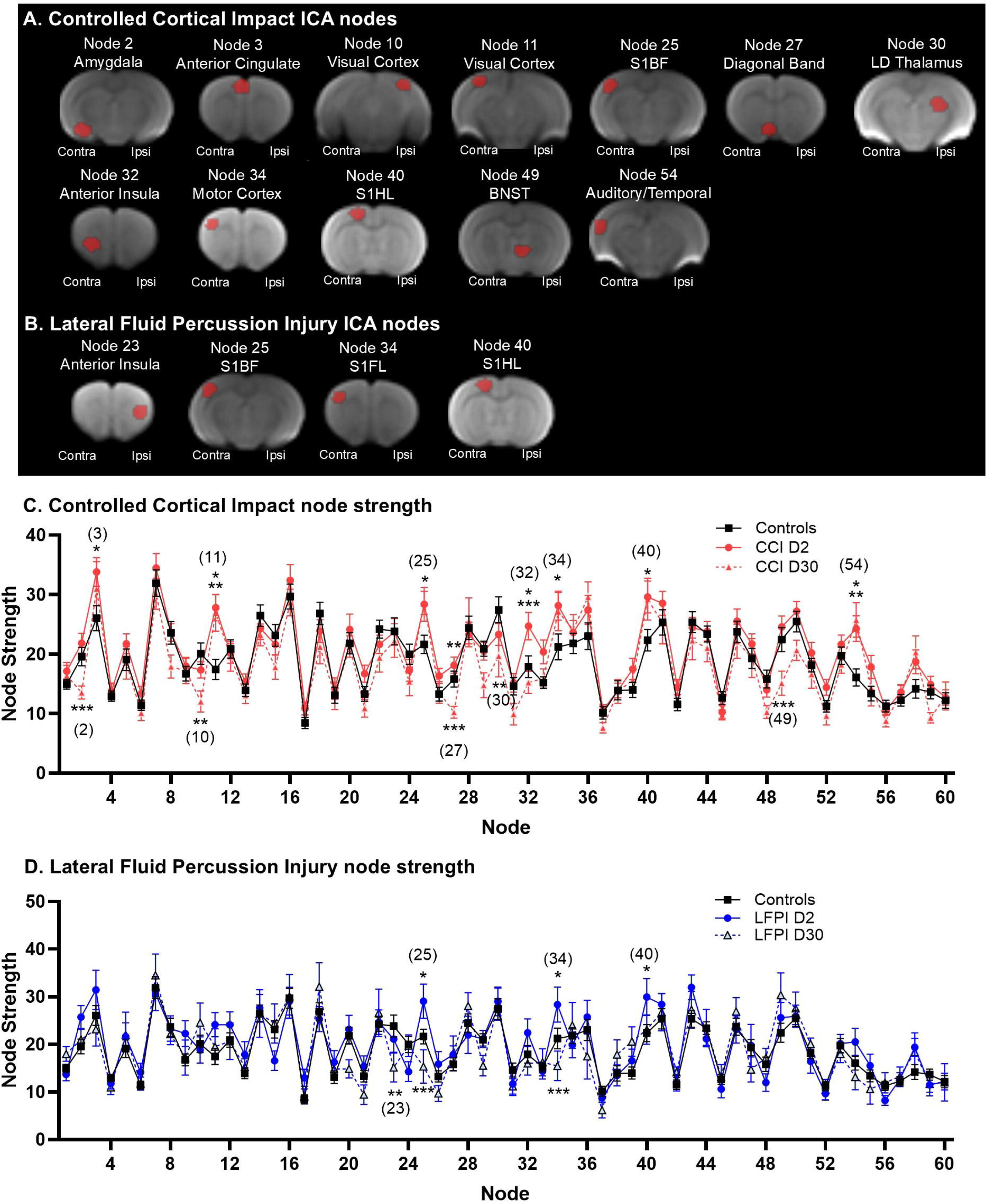
Distinct brain region functional changes in CCI and LFPI. A) Regional distribution of ICA based nodes with significant node strength changes in response to CCI. B) Distribution of nodes with node strength changes in response to LFPI. C) Node strength comparisons between controls, CCI days 2 and 30 in each ICA node. C) Node strength comparisons between controls, LFPI days 2 and 30 in each ICA node. Numbers in parenthesis in each plot in C-D correspond to node numbers in A-B. Two-way ANOVA comparisons followed by Tukey’s multiple comparisons test (*day 2 vs control, ** day 30 vs control, *** day 30 vs day 2). Data shown as mean ± standard error.

We observed differences between LFPI and control rats across several ROIs, albeit in less nodes than observed with CCI (group x ROI interaction: F_118,2773_ = 0.01) (**Fig. 3D**). LFPI increased node strength in contralateral S1BF (p = 0.03, CCI day 2 vs control), contralateral S1FL (p = 0.04, CCI day 2 vs control), and contralateral hindlimb cortex (p = 0.02, CCI day 2 vs control) on day 2. LFPI reduced node strength in ipsilateral anterior insular cortex (p = 0.03, CCI day 30 vs control), S1BF on day 30 (p = 0.001, CCI day 30 vs control), ipsilateral S1BF (p = 0.001, CCI day 30 vs day 2) and contralateral S1FL (p = 0.003, CCI day 30 vs day 2) on day 30.

### CCI and LFPI produce overlapping structural changes on day 2 but only CCI induced structural changes persist through day 30 post-injury

T2 anatomical scans were analyzed for structural differences on day 2 and 30 post-injury. Results are shown on **Fig. 4**. CCI and LFPI produced overlapping structural changes in both contralateral and ipsilateral brain areas on day 2, relative to controls. This is demonstrated as a greater log normalized Jacobian values averaged across the statistically significant brain regions, which included contralateral sensorimotor cortex, both ipsilateral and contralateral hippocampus, ipsilateral striatum, ipsilateral thalamus, ipsilateral and contralateral inferior colliculus and bilateral regions of the cerebellum in CCI and LFPI groups on day 2. Despite the substantial structural differences on day 2 in both CCI and LFPI groups, only the former showed structural differences persisting through day 30. The persistent CCI-induced structural changes were observed in contralateral corpus callosum overlying the dorsal hippocampus, midline areas of the hypothalamus and medial septum, contralateral nucleus accumbens, contralateral diagonal band area, contralateral BNST, midline region of paraventricular thalamic nucleus, contralateral inferior colliculus, periaqueductal grey (PAG), and contralateral subregions of the cerebellum and midbrain (**Fig. 4B**).

**Figure 4.**
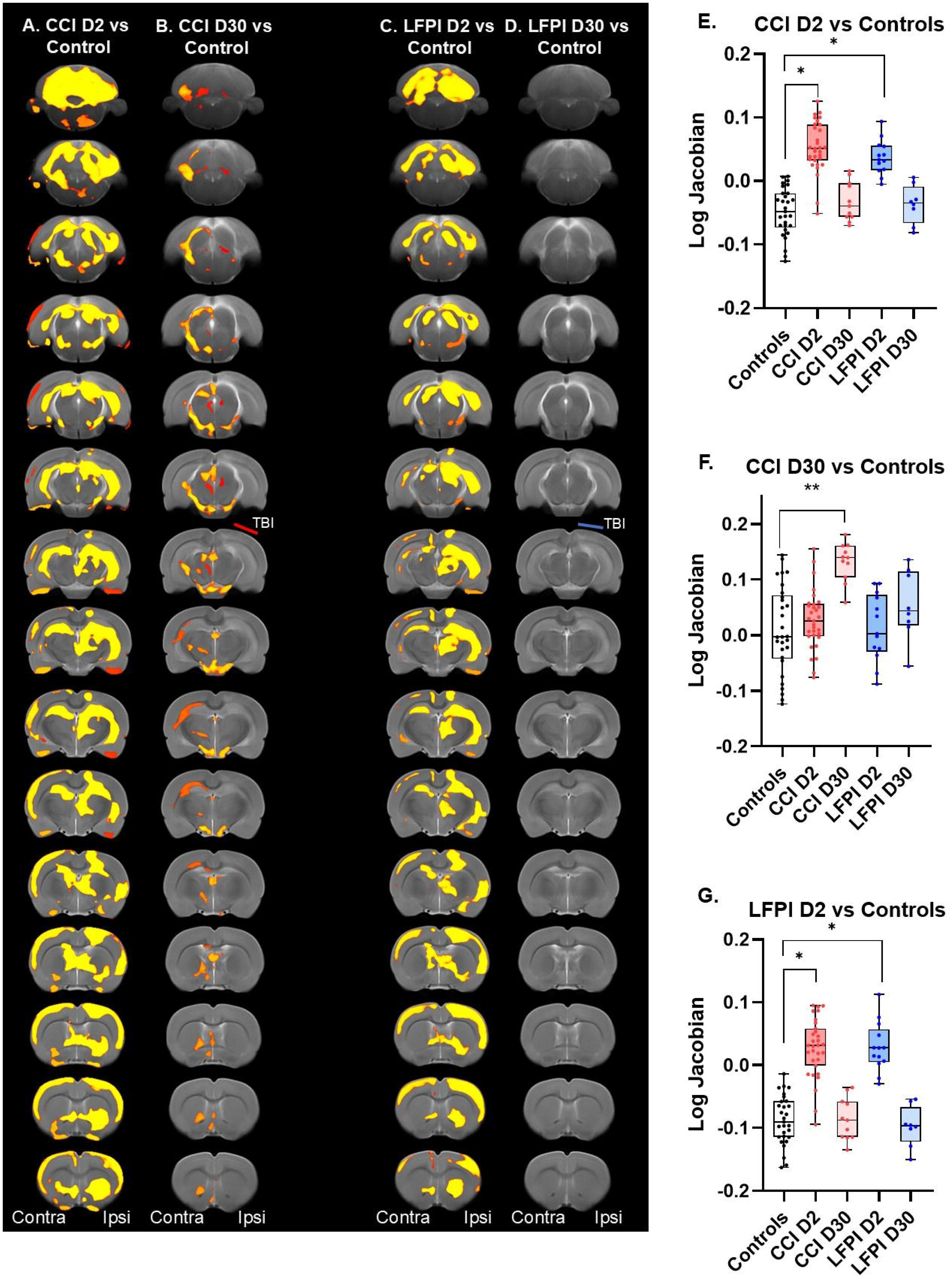
Differential recovery of whole brain structural changes produced by CCI and LFPI. **Maps** on left show areas with significant structural differences between CCI and control on day 2 (A), between CCI and control on day 30 (B), between LFPI and control on day 2 (C) and no differences observed on day 30 following LFPI. E-G) Log Jacobian values for significantly different brain areas highlighted in maps in A-C. The approximate craniotomy/impact site (red bars) for CCI and the craniotomy site for LFPI (blue bars) are shown. One way analysis of variance (ANOVA) with Tukey’s multiple comparisons test (*p<0.05, **p<0.01). Data shown as median and range with overlaid individual data points.

### Distinct White Matter and Cortical Diffusion Changes and Differential Recovery in CCI and LFPI Models

**Fig. 5** shows representative DWI and T2 scans collected in a control, CCI and LFPI rats at 11.1 Tesla, illustrating the slice range where the epicenter was generally located on each of these maps. We analyzed FA, MD, AD, and RD in ipsilateral and contralateral white matter (WM) and cortex. The ROIs included the cortical TBI epicenter and its corresponding section of corpus callosum WM tissue.

**Figure 5.**
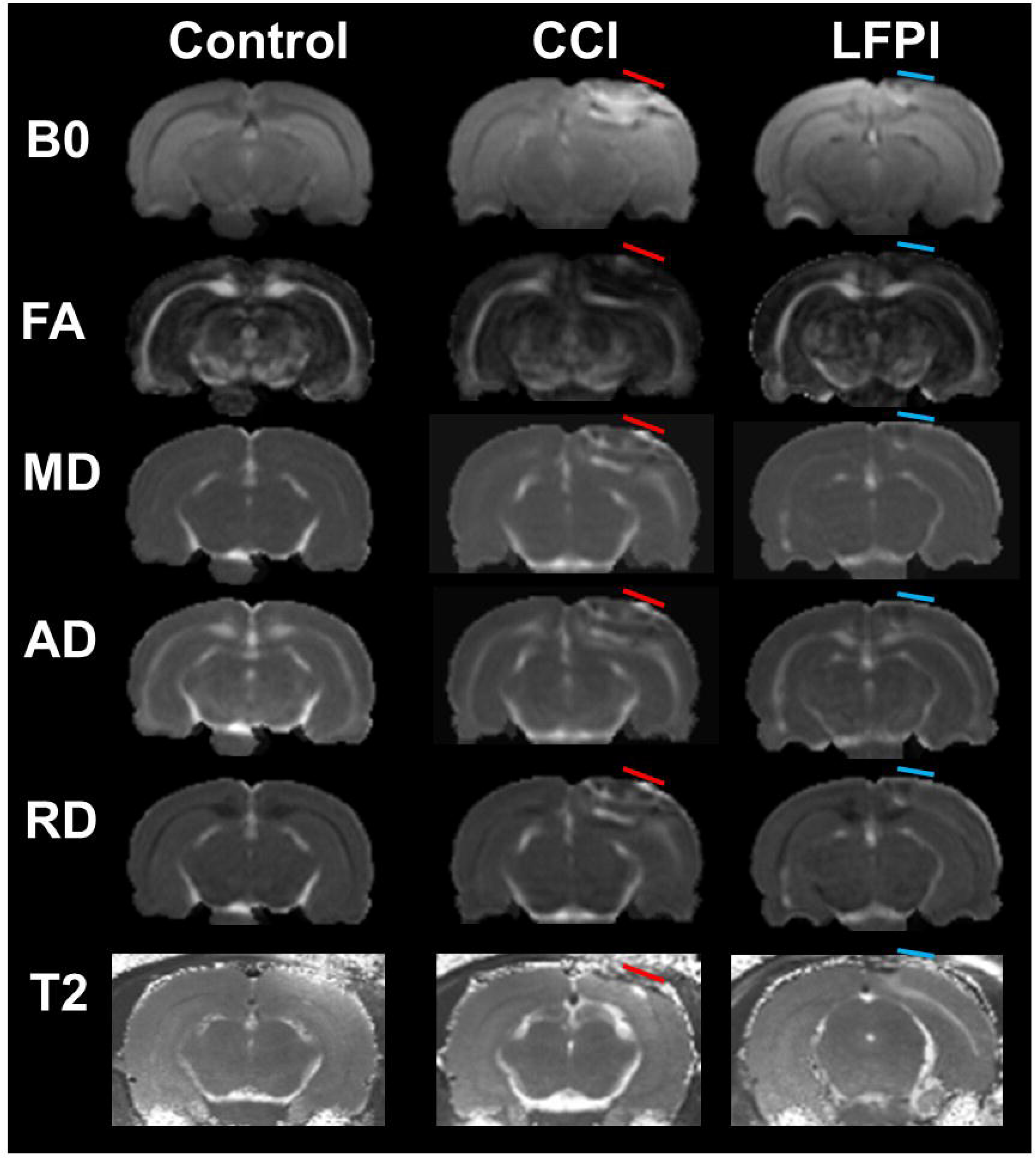
Representative diffusion tensor and T2 maps of a control, CCI and LFPI rats. Shown are a B0 image, fractional anisotropy (FA) grey scale map and mean, axial and radial diffusivity maps (MD, AD, and RD, respectively), and a representative T2 maps of a control animal. Right side is the injury side. The approximate craniotomy/impact site (red bars) for CCI and the craniotomy site for LFPI (blue bars) are shown.

For fractional anisotropy (FA), CCI rats had reduced FA in ipsilateral WM (F_4,85_ = 22.5, p < 0.0001; all post hoc Tukey test comparisons are shown in **Fig. 6**), contralateral WM (F_4,85_ = 12.3, p < 0.0001), and ipsilateral cortex (F_4,85_=4.5, p = 0.002) on day 2 post-injury (**Fig. 6A**). FA returned to control levels in these brain tissues by day 30 post-CCI. LFPI rats had reduced FA in ipsilateral WM on day 2 post-TBI and ipsilateral cortex on day 30 (**Fig. 6A**). No differences in FA between controls and either CCI or LFPI rats were observed in contralateral cortex on days 2 and 30 (**Fig. 6A**).

**Figure 6.**
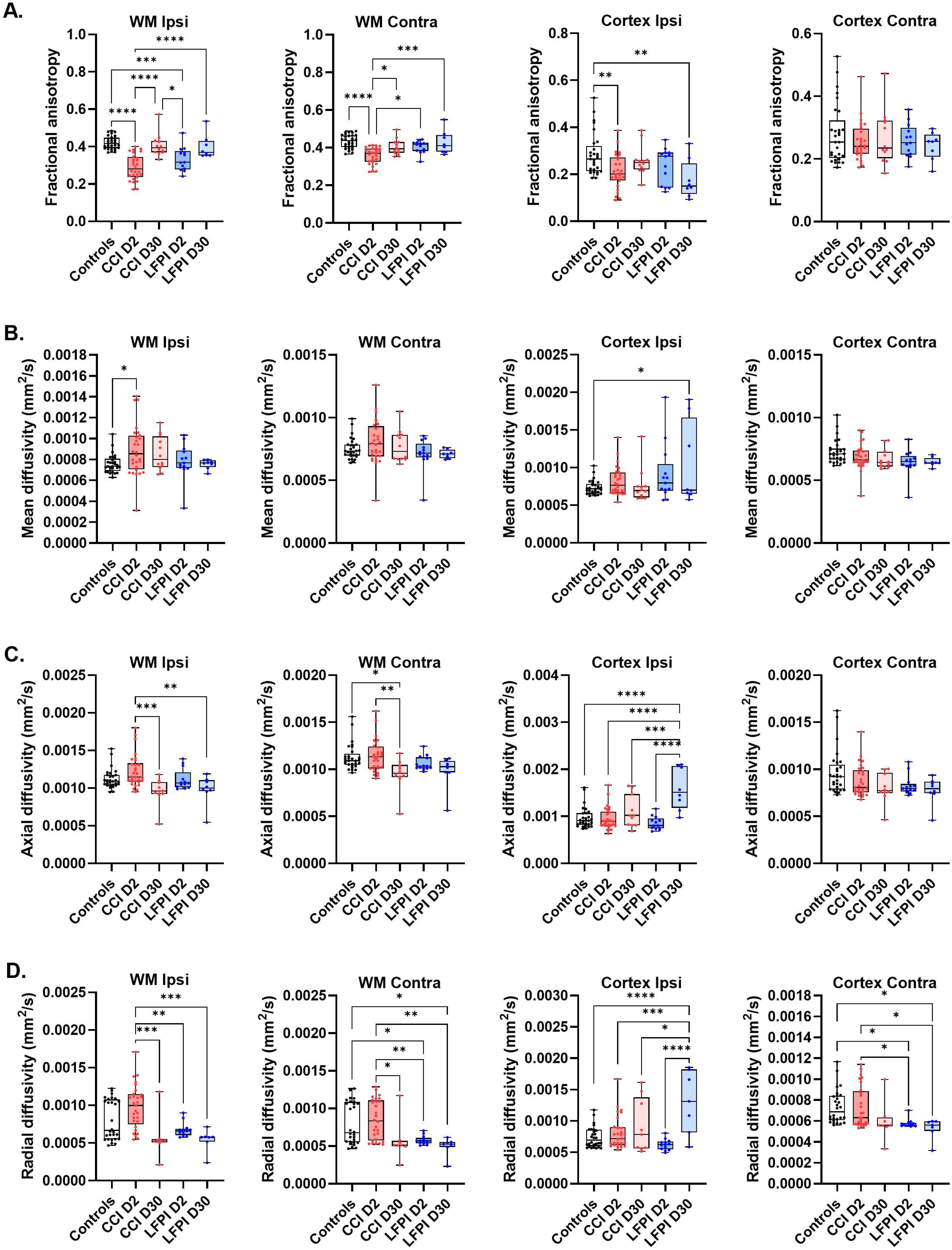
White matter (WM) and cortical tissue diffusion changes at 2 and 30 days following CCI and LFPI. A-D) FA, MD, AD, and RD in ipsi and contralateral WM and cortex. Data are shown as median and interquartile range with overlaid individual data points. and asterisks indicate significant differences at p<0.05 (ANOVA followed by Tukey’s multiple comparison post hoc test; *p<0.05, **p<0.01, ***p<0.001; ****p<0.0001).

Mean diffusivity (MD) in ipsilateral WM was greater in CCI compared to controls on day 2, but not day 30 (F_4,85_ = 2.6, p = 0.04). No differences in MD between control and CCI rats were observed in contralateral WM, and ipsilateral and contralateral cortex (**Fig. 6B**). MD increased compared to controls in ipsilateral cortex on day 30 post-TBI (**Fig. 6B**). No differences in MD between control and LFPI rats were observed in WM ROIs (**Fig. 6B**).

Axial diffusivity (AD) in ipsilateral WM was reduced in CCI rats on day 30 relative to CCI on day 2 (F_4,85_ = 6.2, p = 0.0002) and was reduced in contralateral WM relative to CCI day 2 and to control (F_4,85_ = 4.7, p = 0.001). No differences in AD between controls and CCI rats were observed in the cortex (**Fig. 6C**). AD was greater in the ipsilateral cortex on day 30 in the LFPI rats compared to controls (STATS). No differences in AD between control and LFPI rats were observed in contralateral cortex (**Fig. 6C**).

RD in ipsilateral and contralateral WM was reduced in CCI rats on day 30 relative to CCI on day 2 (ipsilateral: F_4,85_ = 8.0, p < 0.0001; contralateral: F_4,85_ = 6.4, p = 0.0002). Compared to controls, LFPI reduced RD in contralateral WM on day 2, increased RD in ipsilateral cortex on day 30, and reduced RD in contralateral cortex on day 2 and day 30 (**Fig. 6D**).

### Distinct White Matter and Cortical T2 Relaxation Changes and Recovery in CCI and LFPI Models

We analyzed tissue T2 values (in ms) in ipsilateral and contralateral WM and cortex. Compared to controls, T2 increased in ipsilateral WM (F_4,55_ = 7.7, p < 0.0001; all Tukey’s post hoc multiple comparison test results are shown in **Fig. 7**), contralateral WM (F_4,55_ = 2.9, p = 0.03) and in ipsilateral cortex (F_4,55_ = 3.1, p = 0.02) of CCI rats on day 2 post-TBI. T2 values in these brain areas of CCI rats recovered to control levels on day 30. No differences in T2 between control and CCI rats were observed in the contralateral cortex. Compared to controls, T2 values in ipsilateral WM and contralateral WM were greater in LFPI rats on day 2. T2 in these WM areas of LFPI rats recovered to control values by day 30 post-injury. No differences in T2 between control and LFPI rats were observed in ipsilateral and contralateral cortex.

**Figure 7.**
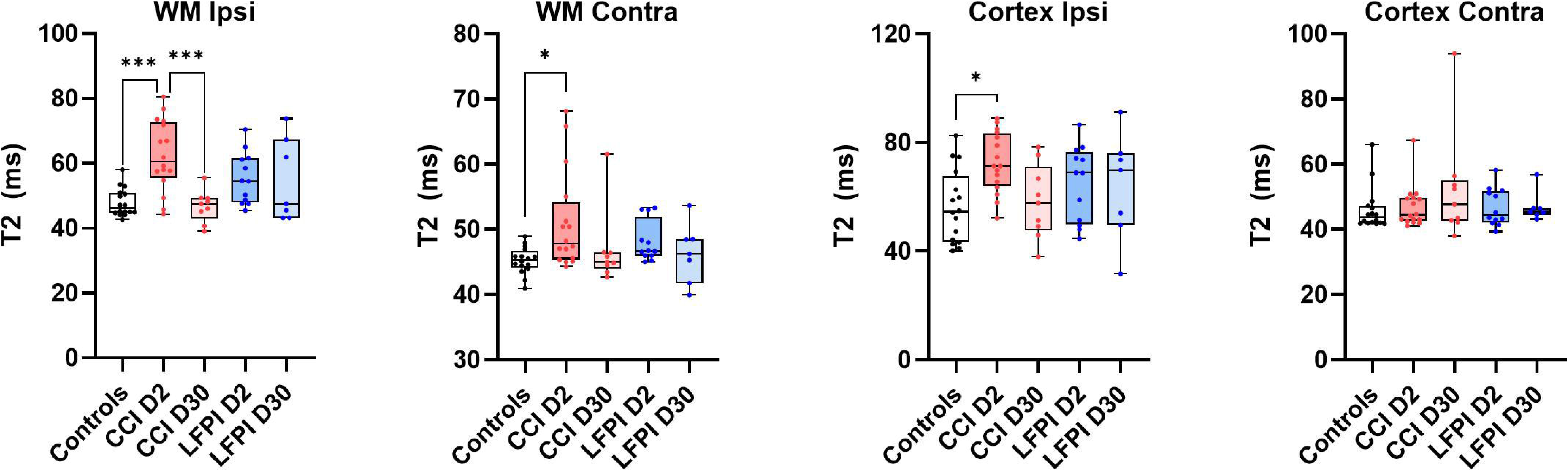
T2 (in ms) in ipsi and contralateral WM and cortex of control, CCI and LFPI exposed rats on days 2 and 30 post-TBI. Data are shown as median and interquartile range with overlaid individual data points. and asterisks indicate significant differences at p<0.05 (ANOVA followed by Tukey’s multiple comparison post hoc test; *p<0.05, **p<0.01, ***p<0.001; ****p<0.0001).

## Discussion

This study demonstrates key similarities and differences in cortical functional connectivity between the controlled cortical impact (CCI) and lateral fluid percussive injury (LFPI) models of traumatic brain injury (TBI). Both models showed significantly increased contralateral intra-cortical connectivity at 2 days post-injury, localized to similar medial and lateral cortical areas. However, this increase persisted until day 30 in CCI rats but not in LFPI rats. These findings suggest ongoing neuroadaptive processes in CCI that may not be as pronounced or sustained in LFPI. While we did not assess excitatory-inhibitory dynamics directly, prior studies hypothesize that functional recovery may involve rebalancing these neuronal processes^48^.

LFPI and CCI are distinct TBI models with unique injury mechanisms: LFPI induces mixed focal and diffuse injuries, often affecting subcortical and contralateral structures, while CCI produces controlled, localized cortical damage. Each model exhibits different structural, functional, and inflammatory outcomes, with LFPI causing widespread axonal injury and acute cytokine responses, and CCI leading to localized tissue loss, blood-brain barrier disruption, and persistent neuroinflammation^49–51^.

Consistent with previous reports^15^, functional connectivity reductions in the ipsilateral cortex were observed in both models. Notably, these reductions resolved by day 30 in LFPI rats but persisted in both medial and lateral cortical regions in CCI rats. Affected cortical areas included cross-modal sensory regions and motor cortex, suggesting that comparisons between CCI and LFPI may offer insights into cortical neuronal adaptations critical for functional recovery.

To explore the link between cortical functional changes and tissue microstructure, we analyzed DTI-based diffusivity measures. Both models exhibited alterations in white matter (WM) and cortical FA and diffusivities, though WM changes were more pronounced in CCI and cortical changes in LFPI. This pattern may reflect greater neuronal recovery in LFPI by day 30, potentially through rebalancing neuronal activity in sensorimotor cortices. Axonal remodeling and compensatory mechanisms, which are well-documented in other injury models^52–54^, could explain changes in axial and radial diffusivities. Specifically, increased axial diffusivity and reduced radial diffusivity in the contralateral cortex of LFPI rats suggest enhanced fiber directionality.

T2 relaxometry results showed that elevated T2 values in both ipsilateral and contralateral WM and cortex at day 2 returned to control levels by day 30. While edema at the injury epicenter likely accounts for the initial increase^18^, it is plausible that contralateral regions are also transiently affected by edematous processes. Future studies employing 3D whole-brain histological approaches may help elucidate the mechanisms underlying these observations.

Changes in cortical excitability are a hallmark of TBI and have been implicated in post-traumatic epilepsy (PTE). In both CCI and LFPI models, layer 5 pyramidal neurons have shown increased spontaneous and evoked discharges, with prolonged burst durations during the first two weeks post-CCI^55^. A reduction in spontaneous inhibitory postsynaptic currents (sIPSCs), possibly due to decreased gamma-aminobutyric acid (GABA) release, has also been reported^17^. Increased contralateral cortical connectivity observed in both models may reflect widespread changes in excitability associated with PTE. However, the relationship is nuanced; decreased functional connectivity has been linked to increased interictal epileptic discharges (IEDs) and shorter seizure latencies in some cases^56^.

Mechanistic insights suggest that TBI-induced hyperexcitability may arise from disrupted cross-callosal excitatory inputs, which normally modulate contralateral inhibitory neurons. This disruption could exacerbate overexcitability in the contralateral cortex^57^. Notably, the relationship between neuronal excitability and functional connectivity is nonlinear. For example, inhibition of mitochondrial function reduces both neuronal firing and connectivity, while increased mitochondrial activity can elevate firing without enhancing connectivity^58^. These findings highlight the complexity of neuroadaptations post-TBI.

Functional connectivity changes may also reflect altered neurovascular coupling. TBI-induced blood-brain barrier (BBB) disruption, astrocytosis, synaptic loss, and microvascular damage collectively compromise the neurovascular unit. Early and sustained changes in cerebral blood flow (CBF) have been documented following TBI. Global CBF reductions of up to 85% in the ipsilateral cortex and 49% in the contralateral cortex have been reported within hours of CCI^59^. LFPI-induced CBF alterations are generally milder but still evident at early time points^60^.

Interestingly, long-term studies show partial neurovascular recovery in contralateral cortical regions at one year post-CCI, suggesting that functional connectivity changes in these regions are not solely driven by vascular mechanisms^61^. Instead, they likely involve complex interactions between neuronal activity, structural remodeling, and vascular recovery.

Our DTI analyses focused on cortical and corpus callosum segments near the TBI epicenter and contralateral cortex. Results revealed increased inter-model variability in diffusivity metrics, consistent with prior literature^14,62^. For example, Harris et al. reported persistent ipsilateral WM FA reductions and corresponding diffusivity changes at one and four weeks post-CCI, supported by histological evidence of demyelination^14^. In contrast, FA reductions in callosal WM recovered by day 28 in other studies, though MD changes persisted^62^. This variability underscores the need for standardized DWI/DTI methodologies to enhance cross-laboratory reproducibility.

Increased T2 values observed on day 2 in ipsilateral and contralateral WM and cortex likely reflect acute edema. This has been linked to upregulation of aquaporin-4, particularly in the cortex^18^. Recovery of T2 values by day 30 aligns with previous studies^63^, though some report persistent T2 increases up to day 42^18^. LFPI-induced T2 changes appear to resolve earlier, potentially reflecting faster resolution of edema or differing tissue damage mechanisms^64^.

This study, by design, mainly focuses on neuroimaging biomarkers and neural network analysis. Thus, the current study did not focus on behavioral or physiological outcomes as measures of injury severity. However, as a parallel investigation within the TOP-NT project, we conducted post-TBI blood-based protein biomarker analysis. Temporal blood-based biomarker data have been shown to provide an objective measure of injury severity across different models. For example, serum biomarkers: Neurofilament-L (NFL) is a marker of axonal injury, while Tau is a marker of neuronal injury and neurodegeneration. Together they serve as robust indices for comparing injury severity across experimental models^65,66^. NFL and Tau are also commonly used to assess injury severity and predicting outcome in clinical TBI studies, reinforces this notion^67,68^. Using serial serum samples from the same CCI and LFPI study cohorts, we noted that both CCI and LFPI have comparable peak elevated serum NFL levels within 48 hours post-injury. Importantly CCI has only slightly higher NFL peak levels (16% higher) compared to the LFPI counterpart (results not shown, as data is being prepared for a separate manuscript submission). Thus, these TBI protein biomarker data suggest that injury severity difference is not the primary driver of observed neuroadaptive processes we observed in our MRI findings when comparing the CCI vs. LFPI datasets. In other words, the neuroadaptive processes are likely attributable to distinct pathological and mechanistic effects induced by the two injury models.

Our results highlight the dynamic and model-specific nature of functional and structural changes following TBI. The persistence of functional connectivity alterations in CCI rats suggests prolonged neuroadaptive processes, whereas LFPI rats demonstrate a greater degree of recovery by day 30. These findings emphasize the utility of multimodal imaging to capture the heterogeneity of TBI-induced changes, offering valuable insights into mechanisms of recovery and potential therapeutic targets. Future studies integrating advanced histological and imaging techniques are needed to further elucidate the spatial and temporal dynamics of cortical and white matter reorganization.

## Conclusions

Our findings underscore the value of multimodal magnetic resonance imaging (MRI) in assessing progressive in vivo changes in brain function and tissue microstructure following contusive and skull-penetrating injuries. Functional MRI (fMRI) and diffusion-weighted imaging (DWI) demonstrate potential for distinguishing pathological differences between LFPI and CCI models, providing crucial insights into their distinct neuroadaptive processes. However, comparisons with existing literature highlight the pressing need for standardized methodologies, data harmonization, and robust statistical approaches to improve cross-laboratory reproducibility. Standardization would facilitate more reliable feature extraction and classification, enabling researchers to evaluate and translate preclinical findings into clinically meaningful applications for TBI^69,70^.

## Author Confirmation/Contribution Statement

**Rohan S. Kommireddy:** investigation, writing (original draft; equal), formal analysis. **Shray Mehra:** investigation, writing (original draft; equal), formal analysis. **Marjory Pompilus:** investigation, formal analysis. **Rawad Daniel Arja:** investigation, formal analysis. **Tian Zhu:** investigation. **Zhihui Yang:** conceptualization, investigation, data curation, project administration, formal analysis. **Yueqiang Fu:** investigation. **Jiepei Zhu:** conceptualization, investigation, writing (review editing), supervision, project administration. **Firas Kobeissy:** conceptualization, writing (review and editing), supervision, project administration. **Kevin K.W. Wang:** conceptualization (lead), methodology, validation, resources, writing (review editing), supervision, project administration, funding acquisition (lead). **Marcelo Febo:** conceptualization, resources, writing (review and editing), software (lead), formal analysis (lead), investigation, data curation, visualization, supervision, project administration, funding acquisition.

## Supplemental Figure Legends

**Supplemental Figure 1. Group probabilistic independent components analysis of rat resting state functional magnetic resonance images collected on an 11.1 Tesla MRI scanner.** Twenty components were identified across 92 fMRI datasets. Components were classified according to location of peak Z statistic voxel.

**Supplemental Figure 2.** Estimation of network metrics on randomized versions of matrices shown in Figure 2C-D. Network measures for CCI days 2, 30 and controls calculated at graph density thresholds of 10, 15, and 20%. D) Network measures for LFPI days 2, 30 and controls. Two-way ANOVA comparisons followed by Tukey’s multiple comparisons test (*day 2 vs control, ** day 30 vs control, *** day 30 vs day 2). Data shown as mean ± standard error. The approximate craniotomy/impact site for CCI and LFPI are shown on 3D connectome maps.

