## Supplementary figures and images for "Functional connectivity, tissue microstructure and T2 at 11.1 Tesla distinguishes neuroadaptive differences in two traumatic brain injury models in rats: A Translational Outcomes Project in NeuroTrauma (TOP-NT) UG3 phase study"

### Supplemental Figure 1

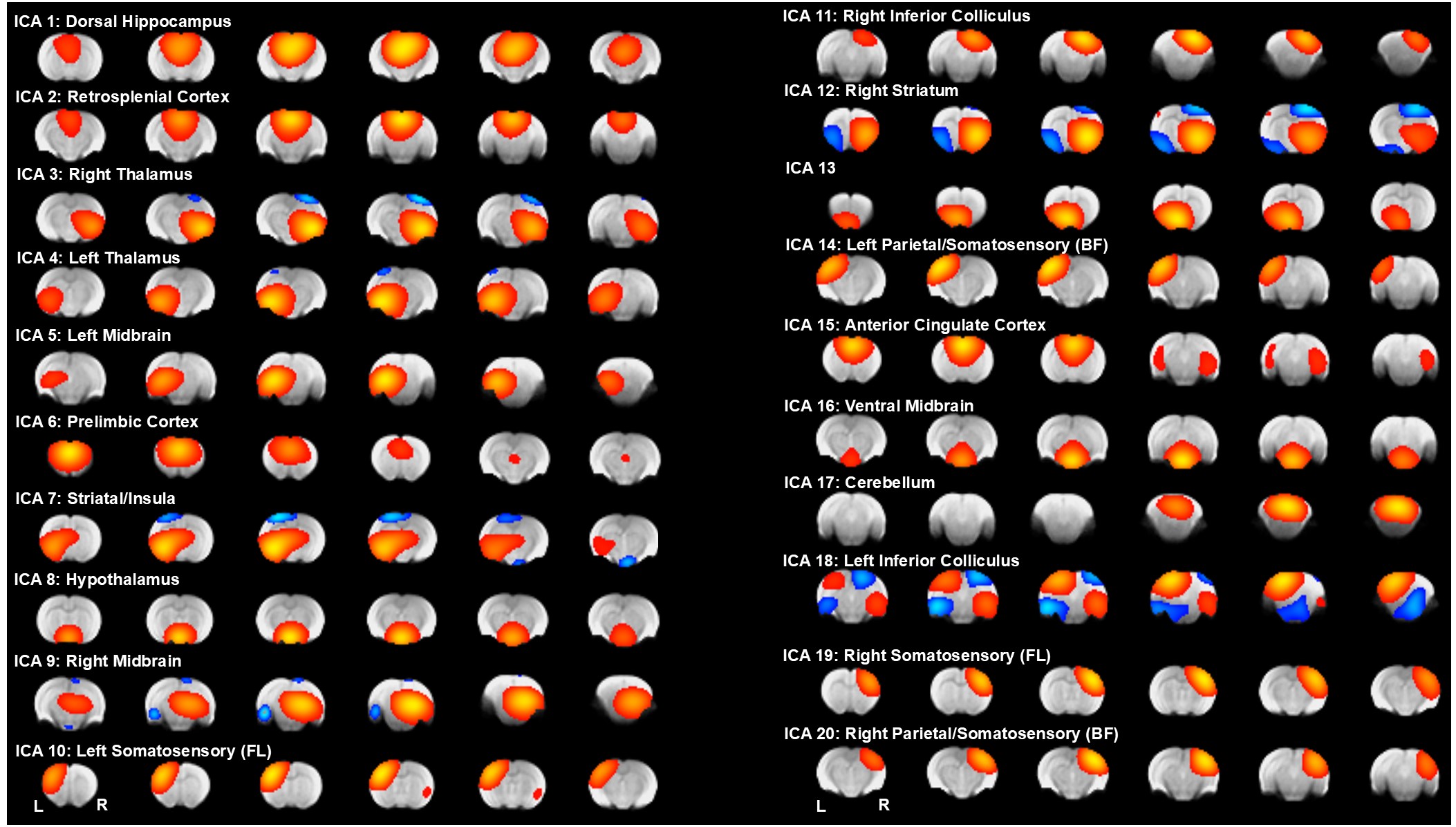

### Supplemental Figure 2

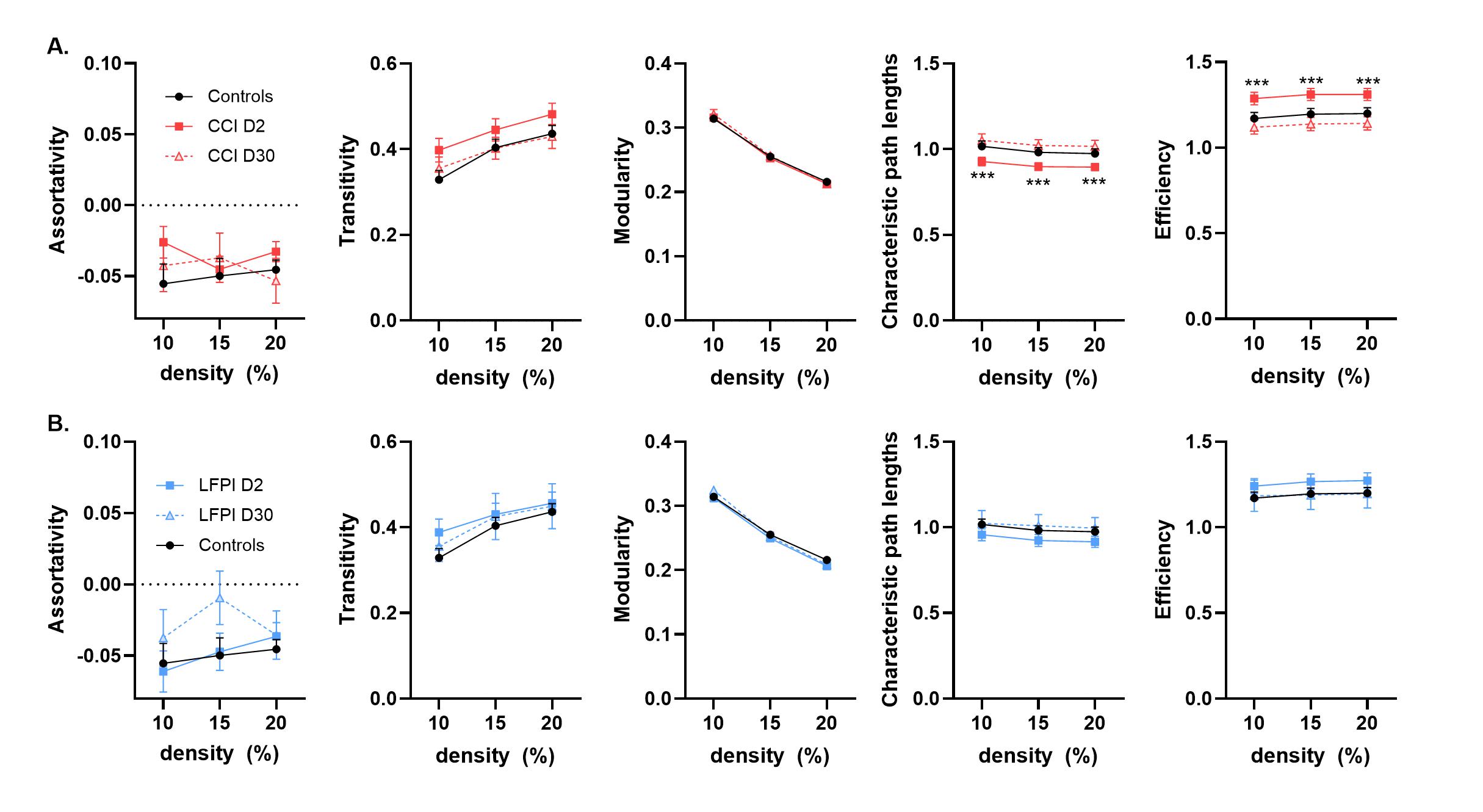
